# ARGContextProfiler: Extracting and Scoring the Genomic Contexts of Antibiotic Resistance Genes using Assembly Graphs

**DOI:** 10.1101/2025.03.24.645075

**Authors:** Nazifa Ahmed Moumi, Shafayat Ahmed, Connor Brown, Amy Pruden, Liqing Zhang

## Abstract

Antibiotic resistance (AR) presents a global health challenge, necessitating an improved understanding of the ecology, evolution, and dissemination of antibiotic resistance genes (ARGs). Several tools, databases, and algorithms are now available to facilitate the identification of ARGs in metagenomic sequencing data; however, direct annotation of short-read data provides limited contextual information. Knowledge of whether an ARG is carried in the chromosome or on a specific mobile genetic element (MGE) is critical to understanding mobility, persistence, and potential for co-selection. Here we developed ARGContextProfiler, a pipeline designed to extract and visualize ARG genomic contexts. By leveraging the assembly graph for genomic neighborhood extraction and validating contexts through read mapping, ARGContextProfiler minimizes chimeric errors that are a common artifact of assembly outputs. Testing on real, synthetic, and semi-synthetic data, including long-read sequencing data from environmental samples, demonstrated that ARGContextProfiler offers superior accuracy, precision, and sensitivity compared to conventional assembly-based methods. ARGContextProfiler thus provides a powerful tool for uncovering the genomic context of ARGs in metagenomic sequencing data, which can be of value to both fundamental and applied research aimed at understanding and stemming the spread of AR. The source code of ARGContextProfiler is publicly available at GitHub.

## 1 INTRODUCTION

Antibiotic resistance (AR) presents a significant global health threat, with an estimated 1.27 million associated deaths globally in 2019 (Naghavi et al., 2024). Horizontal gene transfer (HGT) of antibiotic resistance genes (ARGs) via mobile genetic elements (MGEs) is a fundamental ecological and evolutionary process contributing to the spread of AR across different organisms and environments (Woodford et al., 2011). To develop effective interventions in a “One Health” context, which considers the interconnectedness of human, animal, and environmental health, it is crucial to characterize the evolution and transmission of ARGs within and between associated microbial communities (Aslam et al., 2021; Bustamante et al., 2025).

Methods for characterizing ARGs across One Health-relevant ecosystems; particularly clinical, farm, and broader environmental settings, are needed to identify patterns and trends and inform policy and practice aimed at stemming the spread of AR (Berendonk et al., 2015; Larsson and Flach, 2022). The genomic elements with which ARGs are associated, such as chromosomes, plasmids, or genomic islands, along with neighboring genes, significantly influence their function, regulation, evolution, and the likelihood of undergoing HGT (Aravind, 2000; De and Babu, 2010; Juhas et al., 2009). Therefore, tools that support the systematic exploration of the ARGs in diverse microbial ecosystems, including human and animal microbiomes, agricultural soil, and wastewater, are essential for understanding and mitigating the spread of AR.

Metagenomics provides a means of directly sequencing the collective DNA from a microbial community, offering a more comprehensive view and allowing simultaneous identification and quantification of taxa, ARGs, and other functional genes of interest in a given sample (Hugenholtz et al., 1998; Olsen et al., 1986; Cross et al., 2019; Nogueira and Botelho, 2021). For example, metagenomic analysis of sewage provides a means of profiling ARGs carried across communities served by a particular wastewater treatment plant and holds promise as a monitoring tool that can reveal insightful trends to inform and assess policy interventions aimed at stemming the spread of AR (Bengtsson-Palme et al., 2023; Pruden et al., 2021; Hendriksen et al., 2019). However, genomic context is typically lacking in metagenomic analysis of ARGs. Genomic context refers to the neighboring genetic material present alongside an ARG in a metagenomic sample, which can include additional ARGs, regulatory sequences, and factors influencing gene mobilization. Understanding these genomic neighborhoods is crucial because they drive co-resistance and cross-resistance patterns, ultimately shaping the mechanisms of ARG selection, mobility, and persistence, insights that are vital for designing effective intervention strategies at both local and global scales (Munk et al., 2022) (Fig. 1).

**Figure 1.**
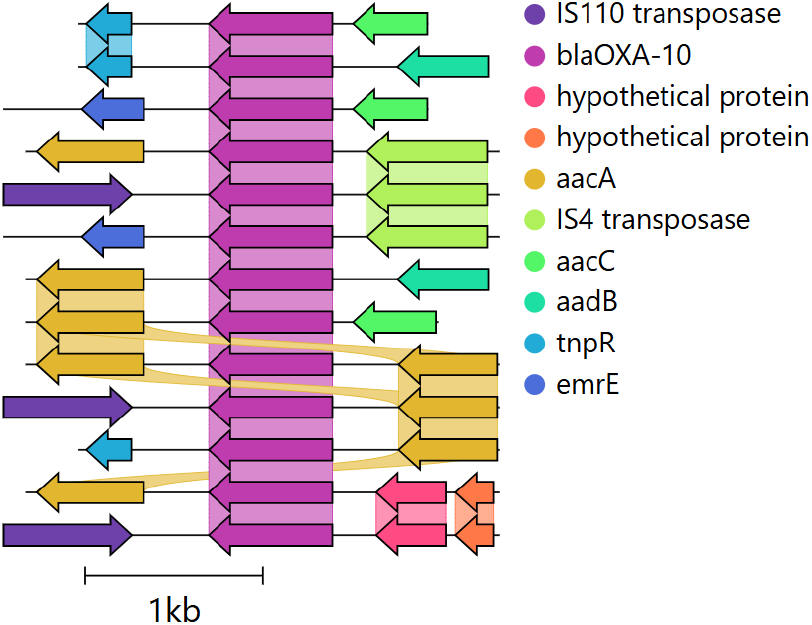
Genomic contexts surrounding the OXA-10 ARG from a hospital sewage sample (ERR1191818), illustrating multiple contexts with MGEs (e.g., transposases) and co-occurring ARGs. The 1,000 bp upstream and downstream regions were identified using ARGContextProfiler, annotated with Prokka, and visualized with Clinker (Seemann, 2014; Gilchrist and Chooi, 2021).

Short-read sequencing remains the dominant approach in metagenomics due to its ability to achieve deep sequencing, along with its cost-effectiveness and high throughput, constituting the bulk of the data available today (Dubey et al., 2022). However, due to the short length of these reads, direct identification of specific ARGs genomic contextual information from these reads is generally infeasible (Arredondo-Alonso et al., 2017; Maguire et al., 2020). One approach to obtain contextual information is to reconstruct the hundreds of millions of short reads into longer stretches, called contigs (Ayling et al., 2020; Zhang et al., 2023). However, assembly can be extremely computationally demanding and confounded by various aspects of the data. For example, highly similar ARG variant sequences that occur in multiple chromosomal and MGE contexts introduce ambiguity that hinders the ability to accurately reconstruct their surrounding sequences (Abramova et al., 2024). Long-read sequencing technologies, like Oxford Nanopore and PacBio single-molecule real-time sequencing, are increasingly used for metagenomics and can provide more comprehensive contextual profiling of the ARGs (Yorki et al., 2023). However, these technologies have limitations, including higher error rates, and, lower throughput, restricting their widespread adoption in metagenomic studies.

Many tools have been developed to assemble short-read sequencing data from metagenomic samples, most of which use variants of de Bruijn graphs approach to handle large amounts of data in an efficient way (Zhang et al., 2023). Subsequently, the tools traverse these graphs and identify the most probable path representing a contig. Converting a graph path into a contig is not a trivial task. Because metagenomic data sets typically contain an unknown number of species with unknown abundance distributions, related species’ sequences can carry similar sets of k-mers resulting in complex assembly graphs. This is further complicated by conserved repetitive regions.

Assembling conserved regions that can occur in several different genomic contexts typically results in highly complex branched assembly graphs, which makes traversing the graphs difficult. This is generally solved by splitting the graph into multiple short contigs. For metagenomic analysis targeting ARGs, this means that sometimes all contextual information regarding the taxonomic origin or mobility of a gene will be lost (Bengtsson-Palme et al., 2017). ARGs constitute a type of genomic feature that is particularly likely to be fragmented in metagenomic assemblies, as they are often present in multiple contexts, can be surrounded by various forms of repeat regions, and can exist on plasmids with varying copy numbers (Abramova et al., 2024).

To tackle these challenges, exploring the intermediate assembly graph generated during metagenomic assembly could potentially be a sensitive method for identifying and analyzing key ARGs within microbial communities. This approach of examining the assembly graph before its conversion into linear contigs has shown promise in tasks such as resolving closely-related strains, SNP calling, and rapid gene homology searches in complex metagenomes (Quince et al., 2021; Alipanahi et al., 2020; Rowe and Winn, 2018). For example, graph-based genomic context analysis tools, like MetaCherchant, often adopt a localized assembly strategy (Olekhnovich et al., 2018). This involves identifying reads and k-mers linked to target genes, followed by constructing a local de Bruijn graph to represent the gene’s vicinity. While effective for highlighting the diversity of the query genes, these methods may not fully capture the broader gene neighborhoods. Other techniques construct the entire assembly graph first, then isolate the query neighborhood, either manually through scaffolding or by automated subgraph extraction methods, such as those used in Spacegraphcats (Brown et al., 2020). However, these extracted subgraphs often include numerous potential paths, lacking a straightforward way to distinguish actual genomic neighborhoods from false chimeric paths. This challenge is particularly notable for mobile ARGs, which can exist in multiple genomic contexts and are frequently linked to repetitive sequences that are hard to assemble. Another approach, Sarand, addresses these issues by utilizing homology searches to explore genomic neighborhoods, combined with coverage-based thresholds to filter out chimeric paths (Kafaie et al., 2023). However, this method has limitations, such as its reliance on heuristic-based graph aligners that might overlook valid genomic paths.

Here we introduce ARGContextProfiler as a means of addressing the challenge of precisely extracting and quantifying valid genomic contexts of ARGs from metagenomic data. This pipeline is specifically engineered to derive genomic contexts of ARGs from metagenomic assembly graphs, enabling a comprehensive assessment of their potential association with pathogens and mobility. ARGContextProfiler employs a sequence homology-based method to pinpoint paths in the graph corresponding to a query ARG, encompassing all possible local upstream and downstream regions. It then implements a series of filters corroborating read pair consistency and variations in read coverage to eliminate chimeric neighborhoods. We rigorously tested the pipeline’s efficacy in reconstructing ARG genomic contexts against the traditional metagenomic assembly process. Our validation involved running ARGContextProfiler on highly complex synthetic metagenomic datasets (CAMI) where the source genomes are known (Sczyrba et al., 2017). We also ran the pipeline on a semi-synthetic dataset (an in-silico spiked human fecal metagenomic sample) as well as on reads from wastewater treatment plants (WWTP) and hospital sewage metagenomes, and compared the genomic contexts captured from these samples to those obtained using standard approaches.

## 2 MATERIAL AND METHODS

### 2.1 Pipeline overview

ARGContextProfiler processes paired-end short reads as input, performs quality control of the reads, and uses metaSPAdes to generate assembly graphs (Nurk et al., 2017) (Fig. 2). The query gene(s) are then mapped to the nodes of the assembly graphs and grouped based on their mapped locations. Each individual instance of the query gene is identified by traversing the graph and extracting the path that represents the gene. For each gene instance, the pipeline retrieves neighboring upstream and downstream regions of the gene up to a user-defined length by searching the graph using the gene path as a seed. Finally, the genomic contexts are constructed and outputted, providing a comprehensive view of the gene’s flanking regions.

**Figure 2.**
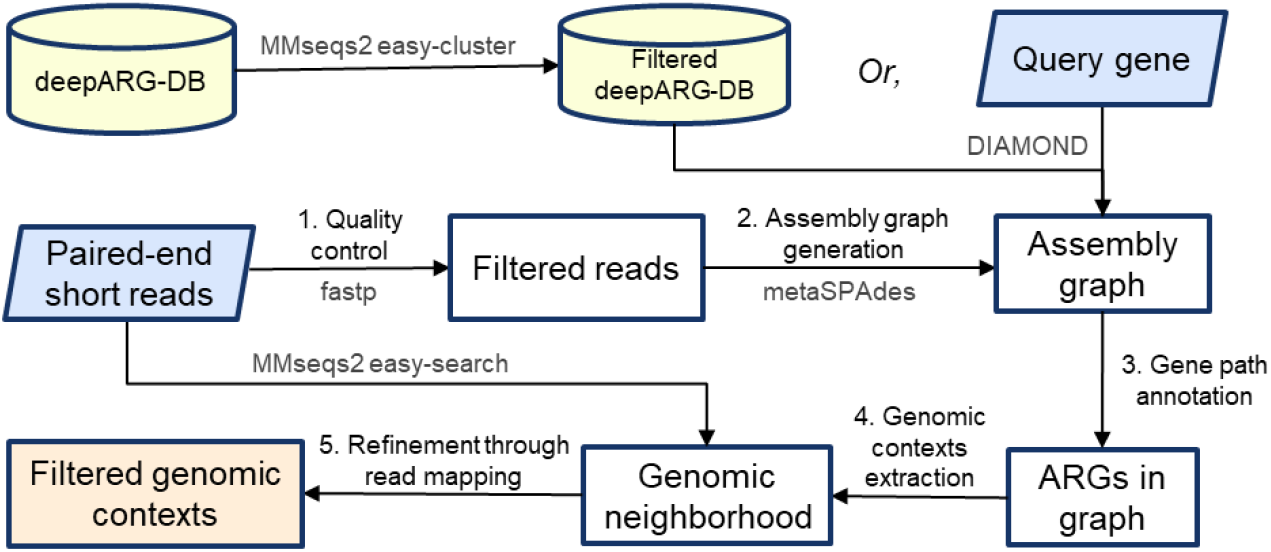
ARGContextProfiler Workflow. The pipeline takes paired-end reads and query gene(s) as input. (1) Reads undergo quality control, and (2) an assembly graph is generated. (3) Query genes are mapped onto the graph, and representative gene paths are identified and annotated. (4) Genomic contexts surrounding each gene path are extracted, and (5) the contexts are filtered through read mapping.

### 2.2 Read Preprocessing and Graph Generation

The pipeline begins by trimming and performing quality control on the raw short reads using fastp (Chen et al., 2018). Following this, an assembly graph is generated using metaSPAdes with default settings and an overlap length of 55 bp. The graph is represented in .fastg format, which provides the sequences of the nodes along with their corresponding overlaps, depicted as edges connecting them. Unlike conventional graphs, the nodes in an assembly graph represent unitigs or DNA segments, each with 3’ and 5’ ends, giving them directional properties. The graph essentially illustrates which ends of neighboring segments are connected, meaning these segments share an overlapping sequence at their prefixes and suffixes.

If query gene(s) are provided, they are mapped to the nodes of the assembly graph using DIAMOND with highly sensitive alignment settings (95% identity) (Buchfink et al., 2015). The mapping is then filtered based on the following condition: if a node has multiple different gene alignments within 100 bp of each other, only the alignment with the lowest e-value is retained. This filtering results in a set of nodes that are mapped to the query gene. If no query gene is provided, the database of ARGs, deepARG-DB is used to map all ARGs to the graph (Arango-Argoty et al., 2018).

### 2.3 Gene Path Annotation

ARGs in the sample are annotated as a sequence of nodes representing a gene path in the assembly graph. The process begins by evaluating the connected components (CC) of the graph to ensure efficient traversal. A CC is defined as a subgraph where any two nodes are connected by a path and no node is connected to nodes outside the component. The coverage of a CC for an ARG is calculated by summing the lengths of all regions that align with the ARG across the entire CC. Only nodes of the CCs that have coverages greater than 60% of the lengths of the ARGs will be selected as “seed nodes” for gene path extraction. Thus paths that do not cover a significant portion of the ARGs will not be traversed, ensuring the quality of the match, and at the same time, significantly speeding up the computation by avoiding traversal of paths that are unlikely to produce quality gene paths. Note that the 60% coverage requirement is set as default and can be changed depending on users’ specific research needs.

Next, for all the seed nodes selected, a recursive depth-first search (DFS) is performed from each seed node to extract potential gene paths from the graph. The traversal is controlled by the following stopping conditions:

#### 1 Sequence gap limit

As nodes are added to the path, if a node does not have homology to the query gene, its entire length is counted toward the cumulative gap. If the cumulative gap of consecutive nodes without homology exceeds a predefined threshold of MAX SEQ GAP, the traversal of that path is terminated. For instance, in Figure 3, path 1 → 2 → 3 is not included as a valid gene path because the gap between nodes 2 and 3 exceeds 1000 bp. Additionally, for nodes that have partial alignment with the query gene, their unaligned portions (either suffix or prefix) also contribute to the cumulative gap. For example, in the path 1 → 2 → 4 → 5 → 6, the gaps are calculated as follows: The unaligned end of node 1, plus the unaligned beginning of node 2 before the aligned region starts; the unaligned end of node 2 after the alignment finishes, plus the entire sequence of node 4 and 5 (since they have no homology with the query gene), plus the unaligned beginning of node 6 before the alignment starts, etc. These individual gaps are summed to determine the total gap size. Since none of the individual gaps or their cumulative sum exceed MAX SEQ GAP, it is considered a valid gene path representing the ARG.

**Figure 3.**
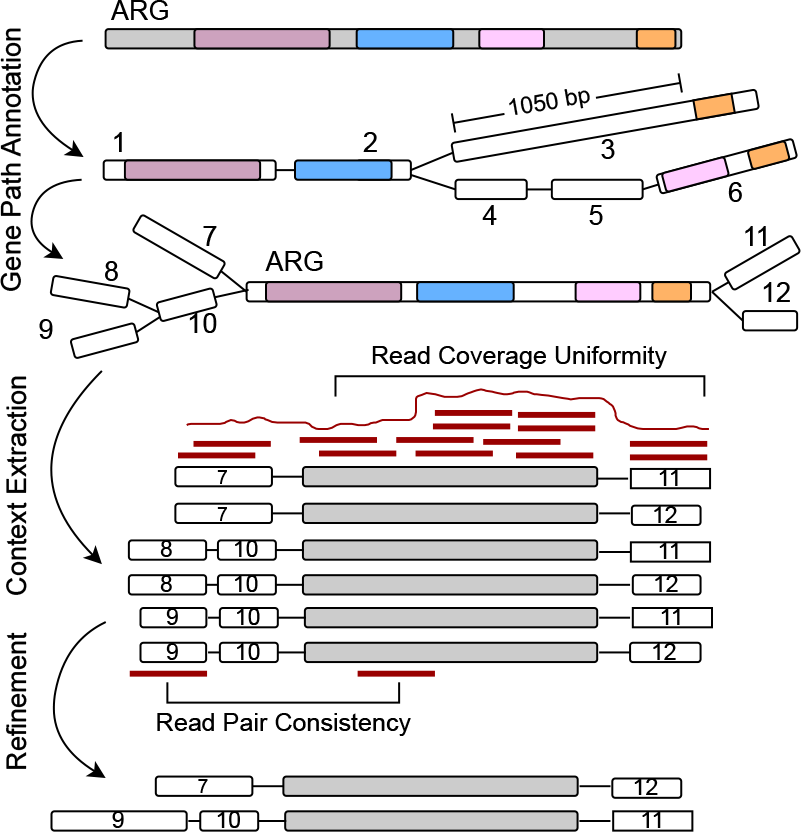
Starting from the alignments on graph nodes for a specific query gene, the graph is explored to extract paths representing the gene. Here, the path 1 → 2 → 3 has a gap of more than 1000 bp of unaligned sequence and is discarded. Only paths with no significant gaps and that cover a substantial portion of the query gene are retained. The neighboring flanking regions are then extracted around the gene path, and chimeric paths are filtered out using read mapping by assessing read coverage uniformity and read pair consistency.

#### 2 Non-homologous node limit

If more than 10 consecutive nodes (i.e., MAX GAP NODE = 10) in the path do not map to the reference gene, the traversal is halted to ensure the path is not overly fragmented. This may also indicate that the end of the gene has been reached within the sequence, making further traversal unnecessary. This condition prevents excessive branching and potential memory overload, leading to a more efficient exploration of gene paths.

After extracting the gene paths, paths that are subsets of others are removed. For each remaining gene path, the gene coverage is calculated by summing the aligned regions from each node along the path. This total alignment length represents the cumulative coverage of the path. If the ratio of this coverage length to the total length of the query gene does not reach a predefined threshold of GENE COV (default is 60%), the path is excluded, as it does not represent a sufficiently significant portion of the gene. The remaining gene paths are clustered using MMseqs2 (95% identity and 95% query coverage) to eliminate redundancy and ensure that the pipeline does not process the same gene multiple times for the downstream contextual analysis (Steinegger and Söding, 2017).

#### 2.4 Genomic Contexts Extraction

To extract the flanking regions, i.e., genomic contexts, of the query gene in the assembly graph, a DFS is used to identify all possible upstream and downstream paths for each representative gene instance. These genomic contexts are explored up to a user-defined threshold (CONTEXT LEN) on both sides of the gene path.

For a given gene path, such as x → y → z, the process begins by exploring upstream paths from the start node x. Since the assembly graph is directed, a path from a+ to b- (indicating the 3’ end of node a connected to the 3’ end of node b, in reverse) implies a reciprocal path from b+ to a-. This directionality is leveraged to extract upstream paths by traversing from the complement side of node x, capturing all possible branches. The traversal is controlled by two stopping conditions:

1. The path reaches the CONTEXT LEN limit.
2. A dead end is reached with no further branching options.

Once all upstream paths are gathered, they are reversed to ensure correct directionality, making the paths terminate at node x instead of starting from x. Similarly, downstream paths are extracted by exploring all possible branches from the end node z of the gene path, also up to the CONTEXT LEN limit. After obtaining both upstream and downstream paths, they are merged with the core ARG path (the gene path x → y → z) to create a complete genomic context for each gene instance. As illustrated in Figure 3, a total of six genomic contexts are recovered by considering all possible combinations of branches from both ends of the query ARG path.

Following this, the combined upstream, downstream, and ARG paths are clustered based on sequence similarity (95% identity and 95% coverage) to remove redundancy and retain only unique genomic contexts. This clustering step ensures that repetitive contexts are eliminated, streamlining the results.

Finally, the resulting genomic contexts are subjected to read mapping-based filters to detect and remove chimeric paths, erroneous combinations of upstream and downstream sequences, that do not originate from the actual genomes in the sample. This filtering step ensures the quality of the extracted genomic contexts surrounding the query gene in the sample.

### 2.5 Refinement through Read Mapping

To eliminate errors, such as inter- and intra-genome misassemblies caused by repetitive genomic regions within the same genome or conserved sequences shared among distinct organisms, we applied a series of read mapping-based filters. These misassemblies often result in erroneous contexts where fragments from different locations are improperly connected, especially in metagenomic samples where ARGs share homology across multiple organisms. Most of these errors manifest as chimeric paths, which are false combinations of upstream and downstream sequences that do not represent a true organism or context.

To detect and remove false-positive chimeric and misassembled contexts, and ensure that the final set of reported genomic contexts accurately reflect the true underlying sequences, we implemented the following two sets of features based on read pair consistency and read coverage uniformity. For more details on the read-based features, refer to Supplementary Section 1.

#### Read pair consistency

For paired-end reads, the insert size (the distance between the left and right mate reads) is assumed to follow a normal distribution *N* (*μ, σ*) (Lai et al., 2022; Wu et al., 2018). The expected insert size *μ* is calculated as the median of all insert sizes, and the standard deviation (*σ*) is estimated by the median absolute deviation. A read is considered proper if its insert size falls within the range [*μ* − 3*σ, μ* + 3*σ*] and its orientation is consistent with its mate. Any read that does not meet these criteria is classified as discordant, which includes three subtypes:

1. Mates mapped to different contexts.
2. Mates with incorrect insert sizes.
3. Mates with inconsistent orientations.

We also classify a read as clipped if it has at least 20 unaligned bases at either end and as supplementary if parts of the read align to different regions of contigs. For each context, the tool calculates the proportion of six read features: proper reads, discordant reads (three subtypes), clipped reads, and supplementary reads. These feature values are analyzed for each context and those outlier contexts having feature values that deviate significantly from the norm are filtered out. Specifically, a context is removed if its mean number of proper reads falls below *μ* − 3*σ* when compared to all contexts associated with the gene. Similarly, contexts are removed if any of the other five features exceed *μ* + 3*σ* compared to the other contexts for the gene. This filtering ensures that only contexts with consistent and well-supported read mapping characteristics are retained.

#### Read coverage uniformity

The read coverage uniformity-based refinement process involves calculating two key features, mean read coverage and normalized coverage deviation, for each context using read mapping data. First, the aligned reads are processed and the coverage for each base across the length of each context is computed. Next, using a sliding window approach, the average coverage and the deviation in coverage (a measure of variation) for each window of length 100bp are calculated. These values are averaged across the entire context, resulting in the mean coverage and normalized deviation. Contexts with high read coverage and low coverage deviation across the length are likely to represent true genomic regions, while those with low coverage or high deviation may indicate potential chimeric or misassembled contexts. Contexts are filtered out based on their mean coverage and normalized coverage deviation in relation to other contexts found for the same gene. A context is removed if its mean read coverage is below *μ* − 3*σ*, where *μ* represents the average coverage of all contexts for the gene, and *σ* is the standard deviation of coverage. This indicates that the context has significantly lower coverage compared to others. Similarly, contexts are excluded if their normalized coverage deviation exceeds *μ* + 3*σ*, where *μ* is the mean deviation and *σ* is the standard deviation of deviation across all contexts for the gene. This threshold ensures that only contexts with consistent, reliable coverage are retained, filtering out those with extreme outlier values.

### 2.6 Data Collection and Preprocessing

The pipeline was evaluated using four datasets:

1. For the fully synthetic metagenomic dataset, we selected the CAMI high dataset from the first CAMI challenge for pipeline evaluation due to its high complexity. With 596 genomes and 478 circular elements, CAMI high provides a challenging test environment and increases the likelihood of encountering multiple genomic contexts for a given ARG, making it ideal for assessing the pipeline’s ability to handle complex metagenomic scenarios.
2. We constructed a semi-synthetic dataset by randomly selecting a metagenomic dataset and spiking it with simulated reads from plasmids containing a known set of ARGs (Abramova et al., 2024) (Fig. 4). The metagenomic dataset, a human stool sample, was downloaded from the Sequence Read Archive (SRA) (SRR9654970) (Leinonen et al., 2010). To select plasmids for spiking, we focused on clinically-relevant and commonly observed ARGs from various classes, including *sul2* (816 bp), *bla*NDM-1 (813 bp), *bla*TEM (861 bp), *aph*(3”)-Ib 3 (804 bp), and *tet*(A) (1200 bp). Protein sequences corresponding to these ARGs were obtained from the Comprehensive Antibiotic Resistance Database (CARD) and used as queries in NCBI BLAST searches to retrieve complete plasmid sequences (Alcock et al., 2023). Only plasmids with *<*98% identity to the query ARG and corresponding to full-length sequences were selected, with five plasmids chosen for each ARG (Abramova et al., 2024). Simulated reads were generated using insilicoseq with the NovaSeq error model (Gourlé et al., 2019). Plasmid read distributions were fine-tuned using an abundance file that assigned higher coverage to smaller plasmids (Abramova et al., 2024). Simulated read sets were generated at 10x coverage levels, with 1x corresponding to the number of reads required to cover the largest plasmid once. To ensure a clean test setup, reads specific only to the human stool sample were first mapped to the selected plasmids, and all matches were removed. The cleaned dataset was then spiked with the simulated plasmid reads.
3. Two real-world metagenomic datasets were used for pipeline evaluation. The first was collected from a local WWTP in February 2021, which also has corresponding Nanopore long reads available. Datasets are available from NCBI (https://dataview.ncbi.nlm.nih.gov/object/PRJNA1083020?reviewer=n59d7m1loo8437kefsh4qm4s7g&archive=biosample) under the accession number PRJNA1083020 (Brown et al., 2024). The second was publicly-available data from hospital sewage, available from the NCBI Sequence Read Archive (SRA) under accession ERR1191818.

**Figure 4.**
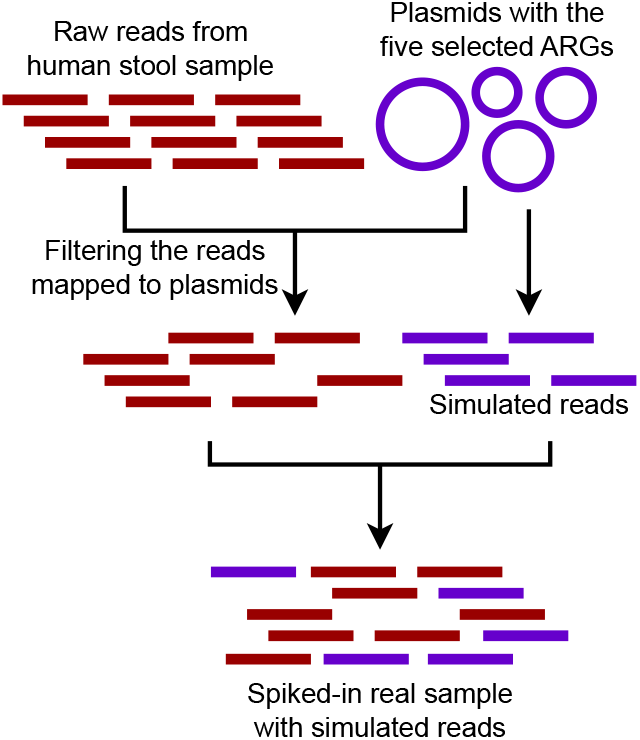
Semi-synthetic dataset composed of a real metagenomic sample spiked with reads generated from a set of plasmids carrying five ARGs.

Reads from all datasets were trimmed, quality-filtered, and decontaminated using fastp and Trimmomatic (Chen et al., 2018; Bolger et al., 2014). Paired-end reads were then used to construct assembly graphs (.fastg) using metaSPAdes (Table 1). The assembly graphs remained unmodified to preserve the neighborhood structure and minimize information loss. For context refinement, reads were mapped to the nodes of the assembly graphs using BWA-MEM, and read mapping-based features were calculated using Samtools (Li, 2013; Li et al., 2009).

**Table 1.** Overview of assembly graph statistics for the four samples generated by metaSPAdes.

| Dataset | Source | Number of Nodes | Number of Edges |
| --- | --- | --- | --- |
| CAMI_high | Synthetic metagenome from a high complexity community with correlated log-normal abundance distributions (596 species and 478 circular elements) (ERS2009087) | 8,909,013 | 471,892 |
| Semi-synthetic | Human stool sample (SRR579292) spiked with plasmids carrying ARGs | 588,356 | 117,142 |
| Hospital Sewage | ERR1191818 | 1,699,140 | 379,796 |
| WWTP | A local WWTP intake sample (PRJNA1083020) | 16,898,439 | 8,908,086 |

For the semi-synthetic, CAMI high, and WWTP datasets, ground truth contexts were derived from the plasmids, source genomes, and corresponding long reads, respectively, by taking 1000 bp (CONTEXT LEN) upstream and downstream of the query gene. These contexts were compared with those derived from ARGContextProfiler and metaSPAdes by alignment using MMseqs2 with the following settings: `--search-type 3 --min-seq-id 0.90 -c 0.90 --cov-mode 2 --max-seqs 1`.

The default values for the parameters were as follows:

GENE COV is set to 0.6, CONTEXT LEN is set to 1000 bp, MAX SEQ GAP is set to 1000 bp, and MAX GAP NODE is set to 10. However, these parameters can be adjusted according to user preference.

### 2.7 Evaluation metrics

Four criteria were used to evaluate the pipeline:

1. **Number of ARGs Retrieved:** This measures the number of ARGs recovered by the pipeline when the exact query ARG is not provided.
2. **Gene Coverage of Extracted Genes:** Gene coverage is defined as the ratio of the recovered length of a gene to its total length in the deepARG-DB. A higher gene coverage indicates that a greater portion of an ARG was successfully reconstructed from fragmented reads.
3. **Precision and Recall:** For the synthetic and semi-synthetic datasets, where the ground truth is known, the contexts derived using ARGContextProfiler were compared with the ground truth. Precision and sensitivity were calculated to assess how many of the reconstructed genomic contexts matched the source genomes (precision) and how many contexts from the source genomes were recovered by the pipeline (sensitivity).
4. **Extracted Context Length:** The lengths of the extracted contexts were also evaluated in order to confirm that the pipeline accurately captured more complete flanking regions. With the context length parameter set to 1000 bp, the expected outcome is an ARG flanked by 1000 bp upstream and 1000 bp downstream, yielding a 1000 bp – ARG – 1000 bp genomic context.

## 3 RESULTS

### 3.1 ARG Detection

We compared the number of ARGs recovered from contigs generated by metaSPAdes and ARGContextProfiler. For the simulated dataset CAMI high, which has ground truth genomes and plasmids available, we assessed the recovered ARGs against the actual number found in the ground truth (Table 2). Notably, nearly half of the ARGs were lost after the assembly process using metaSPAdes while attempting to recover them from the contigs. In contrast, ARGContextProfiler recovered 26 more ARGs than metaSPAdes for this dataset by exploring the graph more thoroughly.

**Table 2.** Number of ARGs recovered by alignment with the deepARG-DB from source genomes, plasmids, or long reads in the CAMI high, semi-synthetic dataset, Hospital sewage, and WWTP sample, respectively; from contigs generated by metaSPAdes and from contexts produced by ARGContextProfiler.

| Dataset | Source | MetaSPAdes | ARGContextProfiler |
| --- | --- | --- | --- |
| CAMI_high | 127 | 61 | 87 |
| Semi-synthetic | 5 | 5 | 5 |
| Hospital Sewage | N/A | 186 | 214 |
| WWTP | 29 | 110 | 166 |

In the semi-synthetic dataset, five ARGs were spiked in silico, all of which were successfully recovered by both pipelines. Each ARG was present in five different plasmids, resulting in high coverage, which facilitated their detection.

For the hospital sewage sample, where the ground truth was unknown, ARGContextProfiler recovered 214 ARGs compared to 186 recovered by metaSPAdes, highlighting its superior ability to navigate this highly complex metagenomic sample.

Finally, for the WWTP sample, the ground truth was based on long reads, which were generated from the same sample as the short reads. However, only 29 ARGs were annotated from the long reads, likely due to low coverage associated with nanopore long-read technology (Hu et al., 2020). Both metaSPAdes and ARGContextProfiler recovered many more ARGs than those found in the long reads. Notably, ARGContextProfiler outperformed metaSPAdes by recovering a total of 56 additional ARGs and captured 27 ARGs confirmed in the long reads compared to metaSPAdes’ 24.

### 3.2 Comparison of Recovered ARG and Genomic Context Lengths

We compared the lengths of recovered ARGs from assembled contigs generated by metaSPAdes and ARGContextProfiler (Fig. 5(a)). The lengths of all recovered ARGs were normalized against their corresponding lengths determined from deepARG-DB. Across all four datasets, ARGContextProfiler consistently recovered longer gene fragments compared to metaSPAdes, indicating that ARGContextProfiler is more effective at retrieving complete ARGs. Genes recovered from metaSPAdes contigs were often fragmented, meaning that only partial gene sequences were retrieved from the contigs.

**Figure 5.**
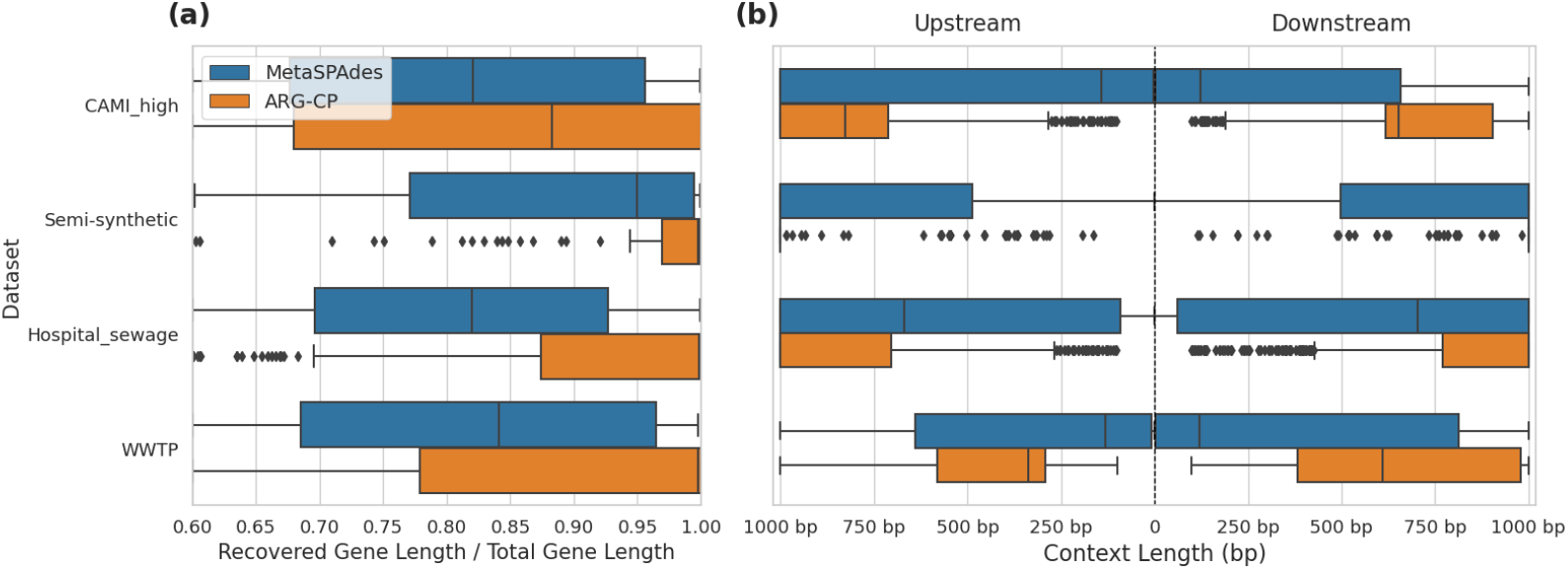
(a) Comparison of recovered gene lengths between metaSPAdes contigs and ARGContextProfiler (ARG-CP) across four datasets: CAMI high, semi-synthetic, hospital sewage, and WWTP. (b) Comparison of the distribution of lengths of extracted ARG contexts in both upstream and downstream directions from metaSPAdes assembly contigs versus contexts generated by ARGContextProfiler across the same datasets.

Additionally, we evaluated the lengths of the extracted genomic contexts in both upstream and downstream directions, further demonstrating the effectiveness of ARGContextProfiler (Fig. 5(b)). In all datasets, ARGContextProfiler recovered longer genomic contexts in both directions compared to the contexts from metaSPAdes contigs.

### 3.3 Comparison of Recovered Contexts in Synthetic, Semi-synthetic, and WWTP Datasets

Using the reference genomes and plasmids for the CAMI high dataset, the reference plasmids for the semi-synthetic dataset, and the corresponding nanopore long reads for the WWTP dataset as ground truth, we evaluated the performance of ARGContextProfiler and compared it to metaSPAdes in extracting 1000 bp upstream and downstream ARG neighborhoods. A relative gene coverage threshold of 60% was applied for both methods.

ARGContextProfiler consistently outperformed metaSPAdes in terms of both sensitivity and precision in recovering genomic contexts in the CAMI high dataset (Fig. 6(a)). While the precision of ARGContextProfiler in identifying valid genomic contexts was comparable to that of metaSPAdes, ARGContextProfiler demonstrated significantly higher sensitivity. On average, ARGContextProfiler achieved a precision of 94.35%, compared to metaSPAdes’ 89.23%. ARGContextProfiler’s average sensitivity was 51.77%, nearly double that of metaSPAdes’ 31.55%. Across nearly all ARGs,

**Figure 6.**
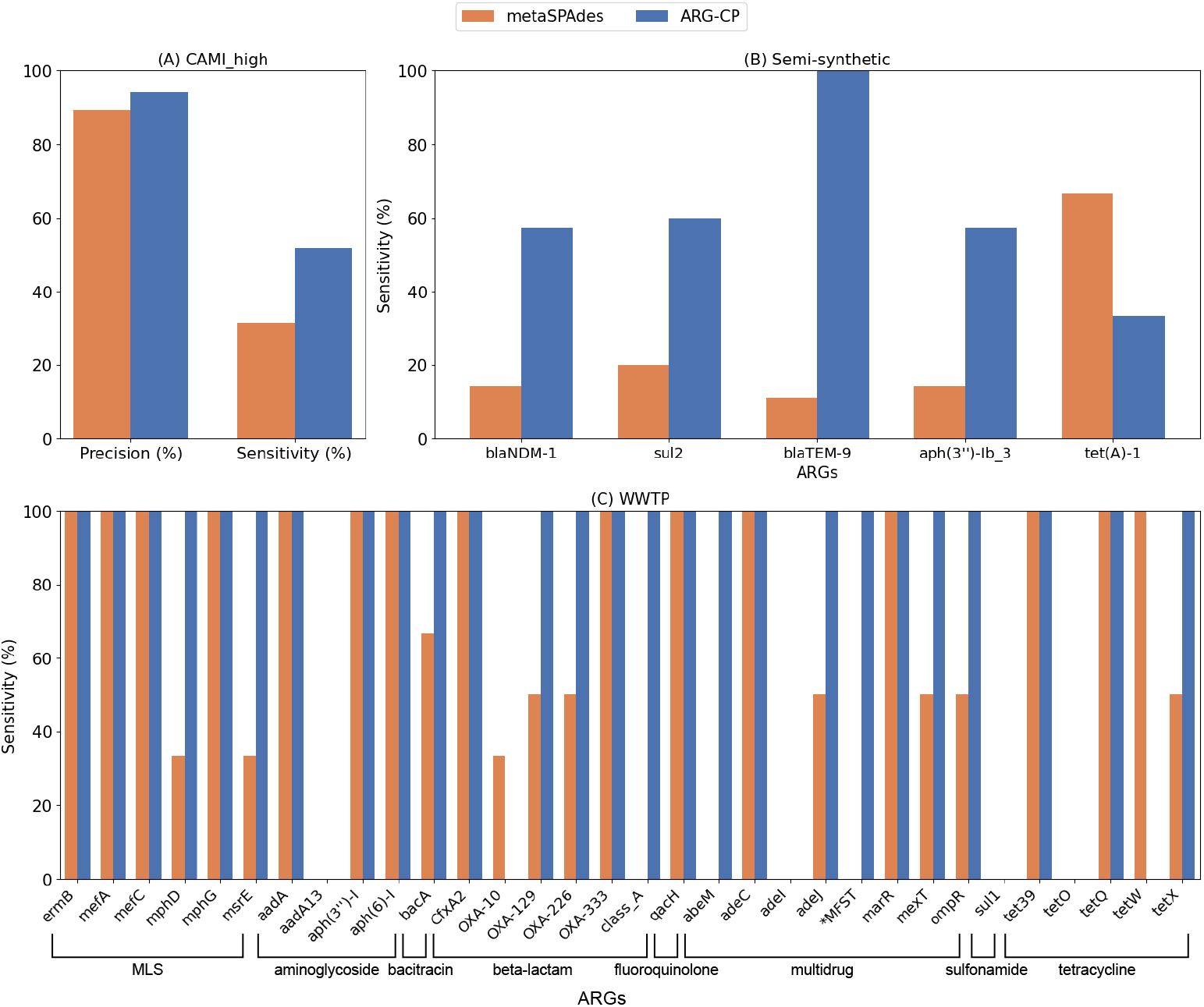
(a) Comparison of average precision and sensitivity between ARGContextProfiler (ARG-CP) and metaSPAdes contigs for the CAMI high complexity dataset, averaged across all ARGs. (b) Sensitivity comparison for the five ARGs in the semi-synthetic dataset between ARGContextProfiler and metaSPAdes. (C) Sensitivity comparison for all ARGs in the WWTP sample, based on alignment with the nanopore long reads from the same sample. *MFST refers to the major facilitator superfamily transporter gene.

ARGContextProfiler captured more valid genomic contexts than metaSPAdes (Supplementary Fig. 1) and minimized the prediction of chimeric contexts (Supplementary Fig. 2).

For the semi-synthetic dataset (Fig. 6(b)), the ground truth genomic contexts for the five ARGs were derived from the 25 source plasmids used to spike the dataset. Both ARGContextProfiler and metaSPAdes achieved perfect precision for all five ARGs, indicating that neither pipeline predicted chimeric contexts. Therefore, precision is not compared in the figure. However, ARGContextProfiler outperformed metaSPAdes in sensitivity, capturing more valid genomic contexts for 4 out of 5 ARGs. On average, ARGContextProfiler achieved 81.25% sensitivity, whereas metaSPAdes averaged 61.46%. As an illustrative case, the *bla*NDM gene was found within three distinct genomic contexts across three plasmids; ARGContextProfiler successfully reconstructed two of these contexts (Fig. 7).

**Figure 7.**
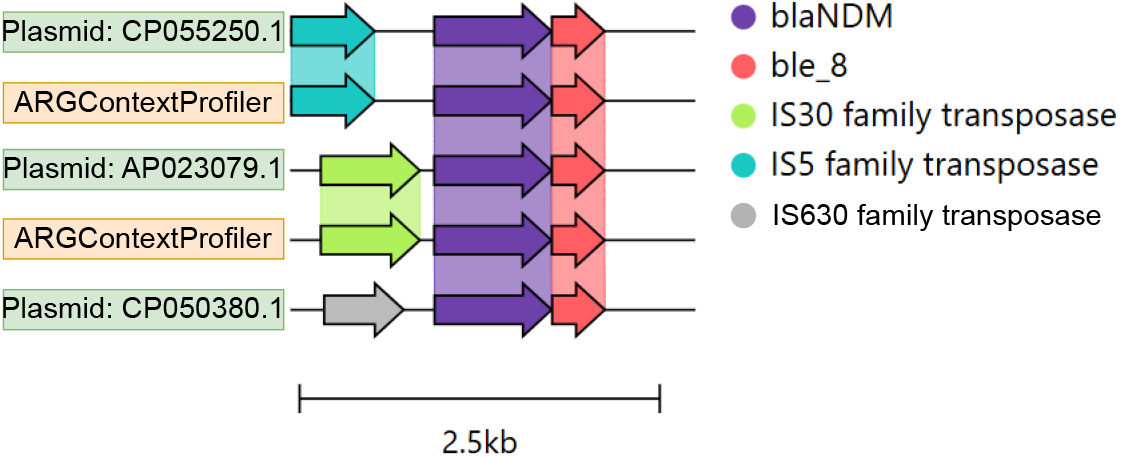
Genomic contexts of the *bla*NDM gene across three plasmids, shown as color-coded arrows. The annotation was performed using Prokka, and the figure was generated using Clinker.

For the WWTP sample, nanopore long reads were also used to validate the recovered contexts. Due to the low-coverage of nanopore long-read technology, many genomic contexts found in both metaSPAdes contigs and ARGContextProfiler predictions were not found in the long reads, making precision difficult to assess. As a result, precision was excluded from the analysis. Instead, we focused on sensitivity, evaluating how many genomic contexts found in the long reads were also captured by each pipeline. As shown in Fig. 6(c), ARGContextProfiler demonstrated superior sensitivity for many ARGs (e.g., *adeM, adeJ, bacA*, class A, major facilitator superfamily transporter, *mexT, mphD, msrE, ompR, tetX*), capturing nearly double the genomic contexts compared to metaSPAdes. However, for two genes (*oxa*-10 and *tetW*), the contigs generated through metaSPAdes captured more genomic contexts (2 and 1, respectively) than ARGContextProfiler. It appears that, due to high sequence similarity (95% identity), ARGContextProfiler annotated *oxa*-10 as *oxa*-129.

## 4 DISCUSSION

### 4.1 Demonstration and Validation of ARGContextProfiler

In this study, we demonstrated that an assembly-graph-based approach to determining the context of ARGs, as implemented by ARGContextProfiler, can greatly improve the quantity and quality of information extracted from metagenomic data sets than relying solely on contigs generated by assemblers. We demonstrated the capabilities of ARGContextProfiler across a variety of datasets, including fully synthetic, semi-synthetic, and real-world metagenomic data derived from WWTP influent and hospital sewage samples. ARGContextProfiler was validated by comparing detected ARGs and their contexts against ground truths, such as in silico spike-ins and long-read sequencing data. In all cases, the assembly graph approach proved superior to metaSPAdes in the detection of ARGs and also in the reconstruction of their genomic contexts. The validity of these contexts was ensured through the use of several read mapping-based features, including read coverage statistics and read pair consistency, which aided in filtering out chimeric or other erroneous genomic contexts.

ARGContextProfiler demonstrated remarkable advantages in terms of the number and quality of contigs derived from metagenomic data, relative to the widely-used assembler metaSPAdes. MetaSPAdes was chosen as a key point of comparison for this study because it is recognized for producing more complete and less fragmented contigs compared to other tools such as Megahit or IDBA-UD, which are known to be vulnerable to the generation of fragmented and incomplete outputs (Abramova et al., 2024; Nurk et al., 2017; Brown et al., 2021; Li et al., 2015; Peng et al., 2012). It is important to mention that, although we validated ARGContextProfiler in part by comparing it to metaSPAdes, it can take assembly graphs as input generated by any other assemblers. We also attempted to compare the analysis of these four large metagenomic datasets using ARGContextProfiler versus other alternatives, such as sarand and spacegraphcats (Kafaie et al., 2023; Brown et al., 2020). However, even when 350 GB of RAM was allocated, both of these tools consistently ran out of memory (OOM). This highlights the substantial memory requirements of currently available methods for large-scale metagenomic datasets, which ARGContextProfiler helps to overcome.

### 4.2 Limitations and Opportunities

There are some limitations to the graph-based assembly approach employed by ARGContextProfiler that should be noted. Firstly, while it recovered more ARGs than metaSPAdes, it is important to note that, like any assembly method, it has a detection limit. As a result, an unknown number of ARGs and their associated genomic contexts may remain undetected in any given metagenomic sample. This is largely due to inherent limitations in the fragmented and incomplete nature of short-read sequencing data, which inevitably introduces some degree of errors during graph construction. When short reads are used to generate an assembly graph, such limitations of the input data may be reflected in the graph’s structure, leading to missing information. Despite these limitations, the graph-based approach still provides a more comprehensive view than directly relying on assembler-generated contigs or short reads alone.

We also highlight that ARGContextProfiler is not limited to the analysis of ARGs. It can be generalized to extract and analyze the genomic context of any queried gene of interest from a metagenomic sample, for example, biocide resistance genes, metal resistance genes, and virulence factors. Users can replace ARG databases with the genes of their interest and apply the pipeline exactly like ARGs. Extending the pipeline to other genes makes it a versatile tool for exploring the genomic neighborhoods of various functionally significant genes. This ability to extract and analyze genomic contexts from metagenomic data opens up new possibilities for understanding gene function and interaction in complex microbial communities, providing valuable insights for applications beyond antimicrobial resistance.

Although our pipeline significantly improves the retrieval of genomic contexts compared to annotations based on final contigs, there remain several opportunities for further enhancement. One potential way to improve sensitivity is to generate a custom assembly graph using the de Bruijn graph constructor ABySS without applying any filtering or bubble removal steps (Simpson et al., 2009). This custom graph, rather than the default SPAdes graph, can help preserve more information and capture more complete genomic contexts (Azizpour et al., 2024). Another option could involve pre-filtering or denoising the assembly graph, although this approach comes with the risk of losing important contextual information. Additionally, an overlap graph, which offers a more precise method than the k-mer-based de Bruijn graph used in most assemblers, could be employed to reduce ambiguity and retain more information (Balvert et al., 2021). However, constructing an overlap graph from complex metagenomic samples is computationally expensive and time-consuming (Li et al., 2012; Rizzi et al., 2019). The use of long-read-based metagenomic graphs, such as those generated from nanopore sequencing, offers another promising avenue (Amarasinghe et al., 2020). These graphs are likely to be more complete, with longer, less fragmented nodes, which could improve both the sensitivity and precision of the pipeline. Moreover, further refinement of genomic contexts can include additional sequence-based filters, such as GC content, protein structure-based features, or the number of open reading frames in a context, all of which would help identify more confident genomic contexts, while filtering out chimeric ones.

## 5 CONCLUSION

ARGContextProfiler was developed as an innovative graph-based approach to profiling the context of ARGs in metagenomic datasets. ARGContextProfiler proved to be a powerful and flexible method for reconstructing genomic neighborhoods of ARG, as was demonstrated over a range of synthetic, semi-synthetic, and real-world datasets, with improved detection, accuracy, and precision relative to metaSPAdes, one of the most widely implemented assemblers currently employed for this purpose. While there are still challenges to overcome, particularly in improving the completeness and sensitivity of the predicted contexts, our results demonstrate that graph-based analysis holds significant potential for advancing the field of metagenomic analysis, particularly when applied to evaluate the context of ARGs, such as their proximity to MGEs. Future developments will focus on enhancing the pipeline’s accuracy and sensitivity, and further investigating the ecological and evolutionary dynamics of ARGs and their genomic contexts, which can potentially uncover new strategies for combating antibiotic resistance.

## CONFLICT OF INTEREST STATEMENT

The authors declare that the research was conducted in the absence of any commercial or financial relationships that could be construed as a potential conflict of interest.

## AUTHOR CONTRIBUTIONS

N.M. conducted the pipeline design, dataset preparation, processing, experimentation, and manuscript writing. The rest of the authors provided continuous support through suggestions and ideas and reviewed the paper.

## ACKNOWLEDGMENTS

This work was partly funded by the National Science Foundation (NSF grants #2319522, #2125798, and #2004751).

## DATA AVAILABILITY STATEMENT

The synthetic, semi-synthetic, hospital sewage, and WWTP datasets can be found under the accession numbers ERS2009087, SRR579292, ERR1191818, and PRJNA1083020 respectively. The source code of ARGContextProfiler is publicly available at https://github.com/NazifaMoumi/ARGContextProfiler.

---

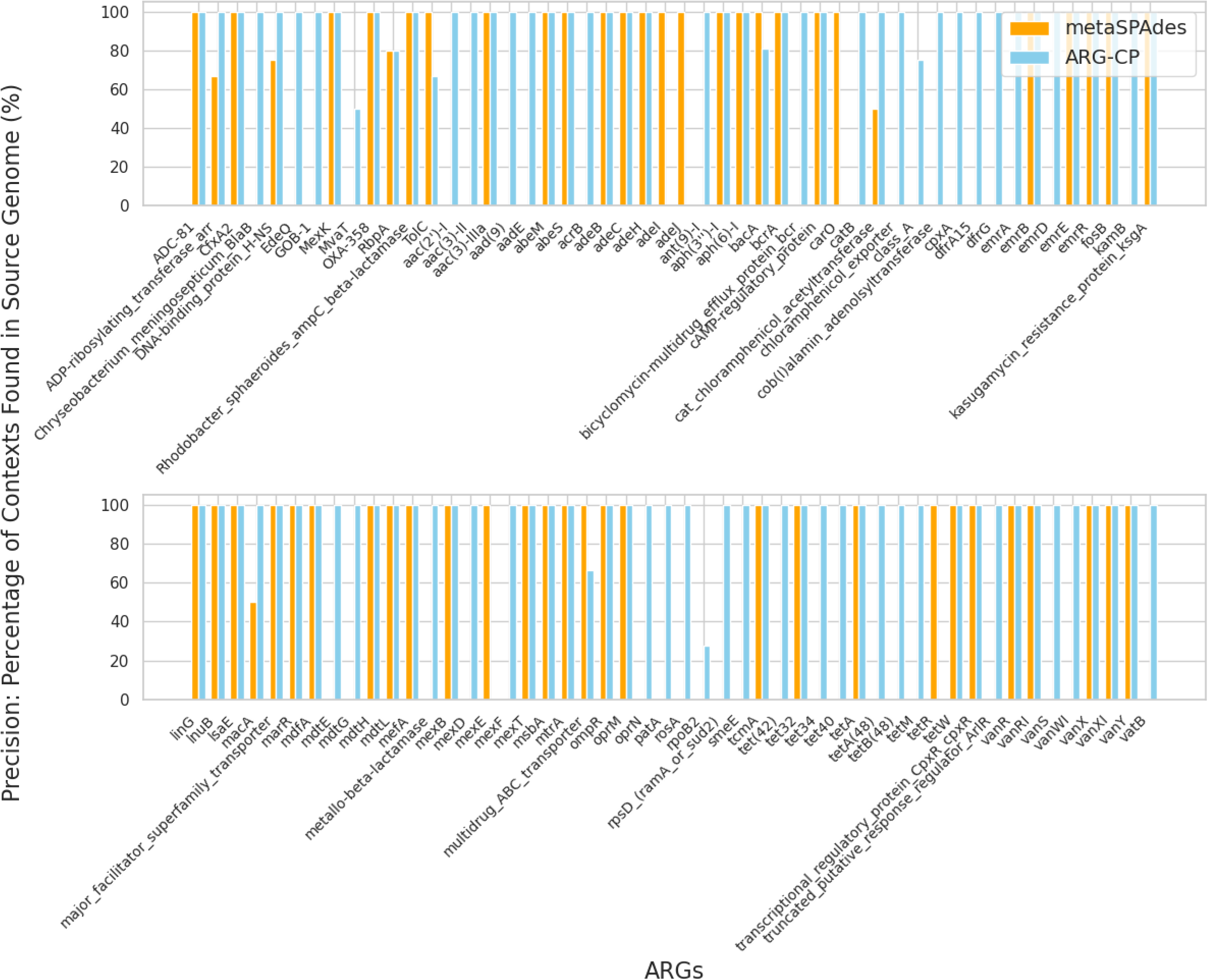

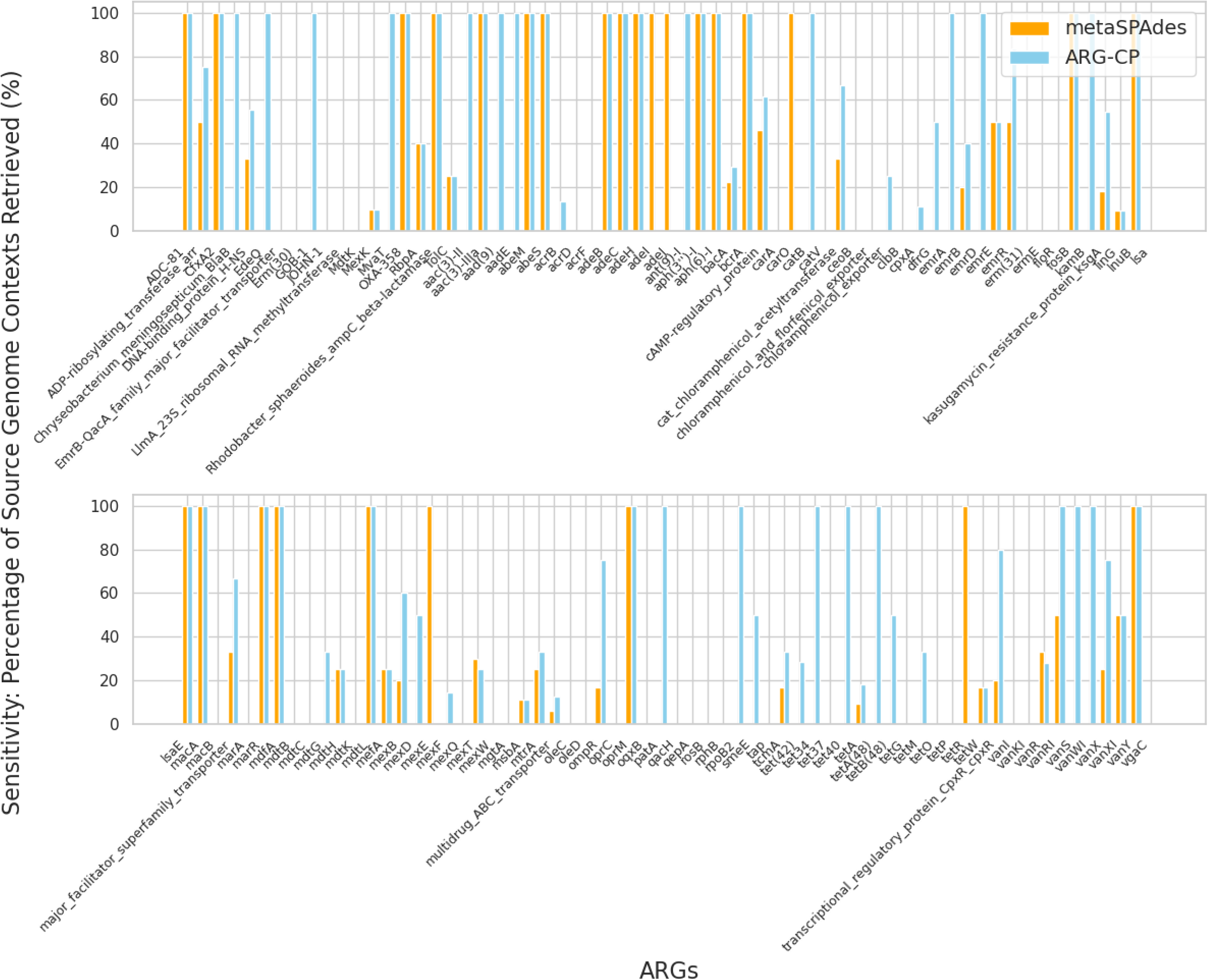

## Notes

### Competing Interest Statement

The authors have declared no competing interest.

