## Supplementary Material for "ARGContextProfiler: Extracting and Scoring the Genomic Contexts of Antibiotic Resistance Genes using Assembly Graphs"

frontiers

### *Supplementary Material*

#### 1 SUPPLEMENTARY SECTION 1

**Read pair consistency-based feature:** Each read is assigned to the following six categories if satisfying the conditions:

**Proper read:**

1. paired-end reads with left and right reads mapping to the same context,
2. the insert size belongs to $[\mu - 3\sigma, \mu + 3\sigma]$,
3. the read pairs have correct orientations.

**Discordant read with a wrong insert size (Type 1 discordant read):**

1. paired-end reads with left and mate reads mapping to the same context,
2. the insert size does not belong to $[\mu - 3\sigma, \mu + 3\sigma]$,
3. the read pairs have correct orientations.

**Discordant read with incorrect orientation (Type 2 discordant read):**

1. paired-end reads with left and mate reads mapping to the same context,
2. The read pairs have incorrect orientations,

**Discordant read with different mapping locations (Type 3 discordant read):**

1. Paired-end reads with left and mate reads mapping to different contexts.

**Clipped read:** the read contains at least 20 unaligned bases at either end of the read.

**Supplementary read:** different parts of the read are aligned to different regions of contigs.

**Read coverage-based features:** Coverage-based statistics including read coverage and fragment coverage are calculated at each base of the assembly by using the information from the input of the BAM file. Read coverage is the number of reads that are mapped over that base at a given base of an assembly. Then read coverage at each position $i$ of context $c$ is further standardized by the following formula, and $L_c$ is the length of the context $c$.

$$\text{Standardized\_read\_coverage}_{i_c} = \frac{\text{read\_coverage}_{i_c}}{\frac{\sum_{j=1}^{l_c} \text{read\_coverage}_{j_c}}{l_c}}.$$

Standardized deviations of read coverage for a given context c are calculated by the formula:

$$\sigma_{\text{read\_coverage}_c} = \sqrt{\frac{\sum_{j=1}^{l_c} \left(\text{standardized\_read\_coverage}_{j_c} - 1\right)^2}{l_c}}.$$

### 2 SUPPLEMENTARY TABLES AND FIGURES

#### 2.1 Figures

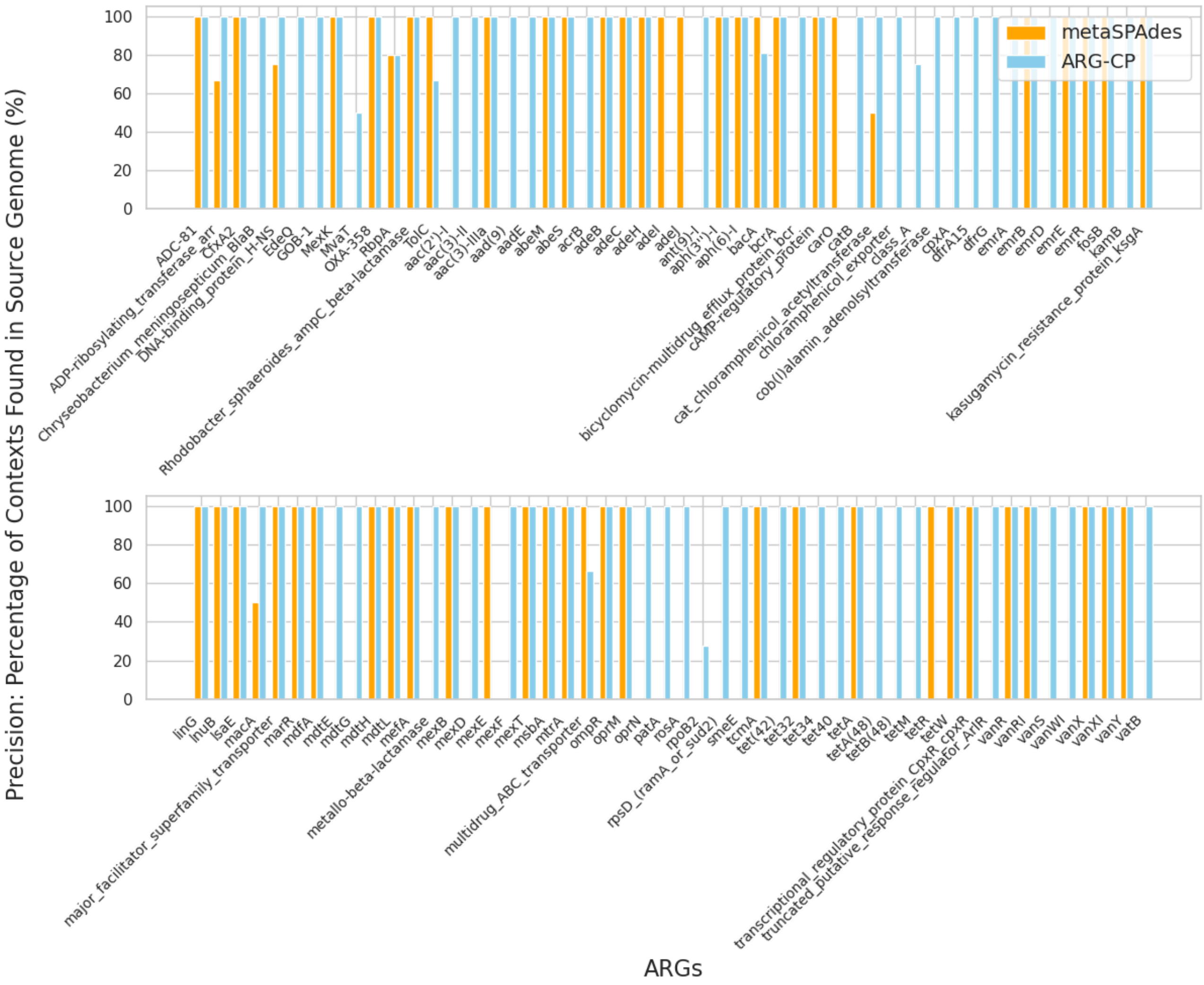


**Figure S1.** Precision comparison for all ARGs in the CAMI_high sample, based on alignment with the source genomes for this sample.

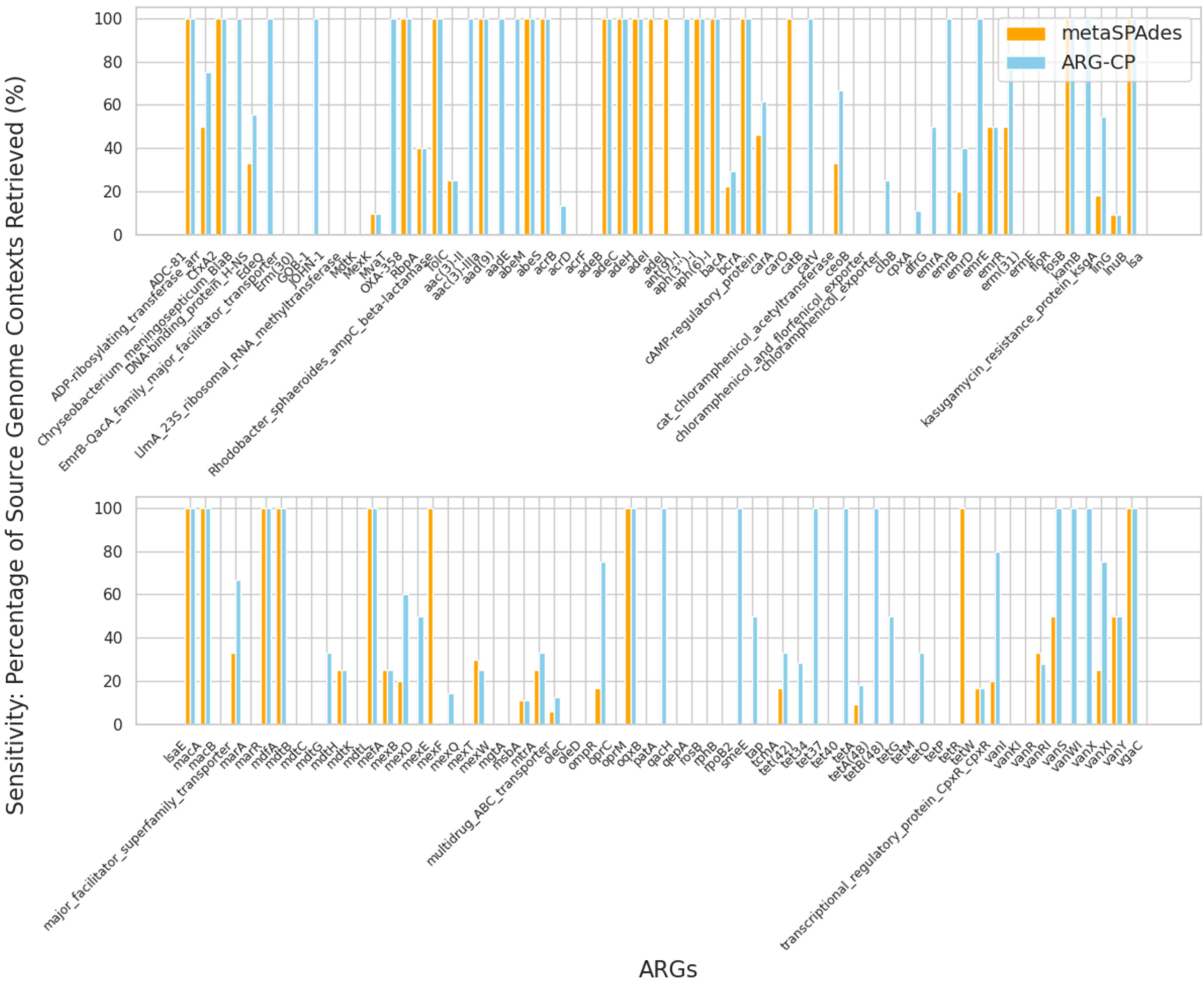


**Figure S2.** Sensitivity comparison for all ARGs in the CAMI_high sample, based on alignment with the source genomes for this sample.
